# Differential bending stiffness of the bacterial flagellar hook under counterclockwise and clockwise rotations

**DOI:** 10.1101/2022.11.08.515729

**Authors:** Xinwen Zhang, Chi Zhang, Rongjing Zhang, Junhua Yuan

## Abstract

The bacterial hook, as a universal joint coupling rotation of the flagellar motor and the filament, is an important component of the flagellum that propels the bacteria to swim. The mechanical properties of the hook are essential for the flagellum to achieve normal functions. In multi-flagellated bacteria such as Escherichia coli, the hook must be compliant so that it can bend for the filaments to form a coherently rotating bundle to generate the thrust when the motor rotates counterclockwise (CCW), yet it also must be rigid so that the bundle can disrupt for the bacteria to tumble to change swimming direction when the motor rotates clockwise (CW). Here, by combining an elastic rod model with high-resolution bead assay to accurately measure the bending stiffness of the hook under CCW or CW rotation *in vivo*, we elucidate how the hook accomplishes this dual functionality: the hook stiffens under CW rotation, with bending stiffness under CW rotation twice as large as that under CCW rotation. This enables a robust run-and-tumble swimming motility for multi-flagellated bacteria.

## Introduction

Most bacteria are able to swim in liquid via rotation of flagella, each consists of a rotary motor, a filament and a hook that couples rotation of the motor and filament [1-7]. The motion states of cells are closely related to the rotational direction of motors. The hook, a flexible universal joint [13], can effectively transmit the rotational states of the motor to the filament [2,14-16]. The flexibility of the hook is crucial for both the turning of swimming direction in monotrichous bacteria [9,17] and flagellar bundling in peritrichous bacteria [18-20].

The hook is a hollow flexible tube with an inner diameter of 3 nm and outer diameter of 18 nm. Approximately 130 copies of FlgE subunits form a 55-nm-length hook in *E. coli* [3,21]. Both the length and flexibility of the hook are optimized for flagellar bundling and steady swimming of bacteria [20,22]. As a torque transmitter component, its mechanical properties were the focus of multiple studies. Block et al measured the torsional rigidity of the hook with optical tweezers [23,24], and found that the hook worked as a linear torsional spring with a torsional rigidity of 1 ×10^8^*dyn* · *cm*^−2^. Flynn et al calculated the Twist/Bend ratio of the hook via a quantized elastic deformational model based on structural information [1], and evaluated a theoretical range of Young’s modulus 10^6^− 10^7^*dyn* · *cm*^−2^ for hooks. The measurement of bending stiffness for the hook in vitro was first accomplished by Sen *et al*. [25]. They evaluated the bending stiffness of the hook on the order of 10^−29^*N* · *m*^2^ by combining electron micrographs of curved hooks and thermal bending. Son et al. found that the bending stiffness of the hook in *Vibrio* was significantly different depending on the load condition, with a value of 2.2 × 10^−25^*N* · *m*^2^ under load of steady swimming and 3.6 × 10^−26^*N* · *m*^2^ for the unloaded state [9]. Nord *et al*. measured the elastic response of a single hook under increasing torsional stress, and found that the bending stiffness of the hook increases in the range of 5 × 10^−26^ −3 × 10^−24^*N* · *m*^2^ as the motor torque increases [26].

Different motor rotational directions (CW or CCW) in *E. coli* correspond to different states of cell movement [27]. When the motor rotates CCW, the hook must be compliant enough so that the filaments can form a coherent bundle at one end of the cell (in a run). When the motor rotates CW, the hook must be stiff enough so that the filament can come out of the bundle and point to a different direction (in a tumble). This raises the following question: Is there any difference in the hook bending stiffness between the two motor rotational directions? A measurement of the hook bending stiffness was attempted recently for CW and CCW-rotating motors [26]. However, a conclusive result was not obtained because of the noise of the measurement.

Here we propose a novel way to measure the hook bending stiffness *in vivo*. Using the bead assay with high spatiotemporal resolution, in which a micrometer-sized latex bead was attached to the shortened filament and the rotation of the bead was monitored, we induced different bending magnitudes of the hook by supplying external fluid flow with different flow speeds. Using a geometric model based on elastic theory, we treated the hook as an elastic rod (instead of approximating it as a linear torsional spring as done previously) and obtained the hook bending stiffness more accurately. We performed measurements with CCW and CW-rotating motors separately, and found that the hook bending stiffness of CW-rotating motors was 1.9 − 2.2 times as large as that of CCW-rotating motors under the same load. Therefore, when the motors rotate CCW, the hooks are soft and promote the bundling of the flagella. In contrast, when the flagella rotate CW, the hook bending stiffness increases, and this promotes the flagella to detach from the bundle and be more perpendicular to the cell surface, thereby helping the cell to change swimming direction.

### Geometric bending model for the hook with bead assays in a flow field

The hook was previously usually approximated as a linear torsional spring with spring constant *EI/L*_*hook*_, where *EI* and *L*_*hook*_ are the bending stiffness and length of the hook, respectively [9,26]. Here, to describe the mechanic properties of the hook more precisely, we treated the hook as an elastic rod with a circular section [28].

Our experiments could be described as the motion of a rotating flagellum marked by a micrometer-sized bead under a horizontal flow field (FIG. 1a). The proximal end of the hook is embedded vertically on the cell surface, while the distal end is connected to a truncated filament stub that is marked by a bead. We treated the hook as an elastic rod with a circular section, while the filament portion was treated as a rigid body due to its much greater stiffness than the hook [1]. Since the force exerted by the flow field on the bead is much greater than that exerted on the hook, the effect of the flow field on the hook itself was neglected here. We can now simplify the model as the bending of an elastic rod that is subjected to a concentrated force f and bending moment *M*_*b*_ = *L*_*stub*_ ×*f* at the free end, where *l*_*stub*_ is the moment arm provided by the filament stub.

**FIG. 1.**
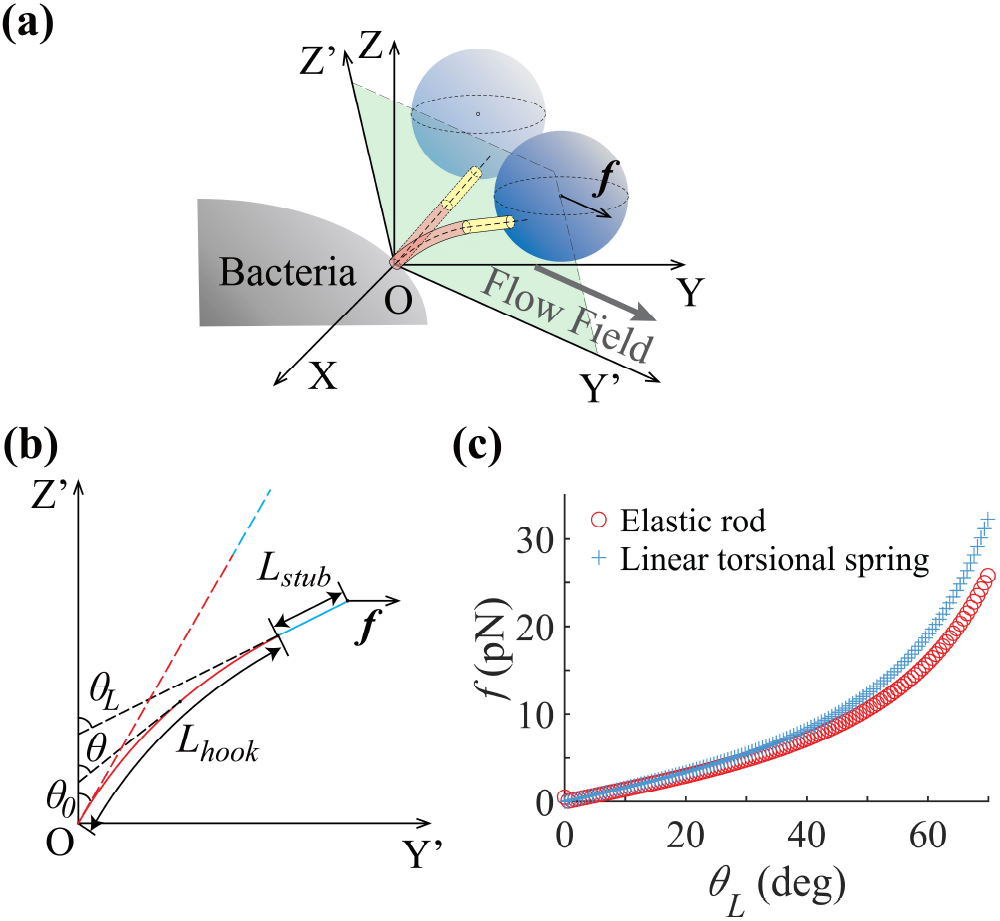
**(a)** Microscopic geometry model of the bead assays in a constant flow field (not to scale). XOY is the image focal plane, and Y’OZ’ is the hook bending plane where OY’ is its intersect with the plane XOY. **(b)** The hook centerline in the bending plane under a concentrated force ***f***. The dashed lines represent the state of the hook (red) and filament stub (blue) before applying the force. The solid lines represent the equilibrium state of the hook (red) and filament stub (blue) under the force. **(c)** Comparison of the relation between the flow field force and *θ*_*L*_ for the elastic rod model (red circles) and linear torsional spring model (blue crosses).

The hook is considered straight along the normal direction of the cell surface in the absence of an external force. Thus, the external force f must be in the bending plane (green shade plane Y’OZ’ in FIG. 1a), and the model is further simplified as a two-dimensional bending problem. As shown in Fig. 1b, an elastic straight rod is embedded at the origin of coordinates O with an angle *θ*_0_ from the Z’-axis. A concentrated force f parallel to the Y’-axis is applied at the free end of the filament stub, and the rod is bent to balance. In equilibrium, the angle between the tangent line of the free end of the rod and the Z’-axis is *θ*_*L*_. The equilibrium differential equations for any element *dl* of the rod can be written as

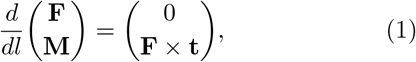

where ***F*** is the resultant vector of stress applied to the cross section of the rod, ***M*** denotes the torque synthesized from stress acting on the cross section of the rod, and *t* is the unit tangent vector of the rod. Thus, we can obtain the equation of bending of the rod:

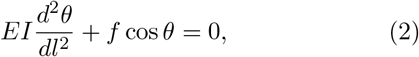

where *E* is the Young’s modulus and *I* denotes the principal moment of inertia for the rod. Therefore, we can compute the bending stiffness *EI* by integrating the above equation (see Supplementary Note 1 for details):

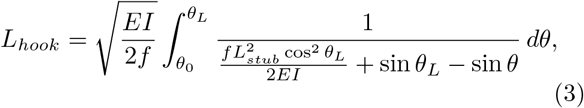

where *θ*_0_ and *θ*_*L*_ denote the angle between the proximal and distal tangents and the Z’-axis, respectively, f is the horizontal force exerted on the bead by a constant flow field, and *L*_*stub*_ is the length of the filament stub, defined as the distance between the hook/filament junction and the bead attachment point.

To compare with the model of linear torsional spring, we used typical values of *EI* = 1 × 10^5^ *pN* · *nm*^2^, *L*_*stub*_ = 200 *nm, L*_*hook*_ = 55 *nm* and *θ*_0_ = 0. The relation between *θ*_*L*_ and *f* from the elastic rod model can be calculated using equation (3). As shown in Fig. 1c, the result from the model of linear torsional spring was drawn with the function *f* (*θ*_*L*_) = *EI/L*_*hook*_ *θ*_*L*_. Apparently the linear approximation can describe the bending of the elastic rod at small angles (e.g., *θ*_*L*_ *<* 30^*o*^), while it significantly underestimates the bending stiffness when the bending angle is large. In reality, the swimming *E. coli* requires the hooks to bend at large angles in order to form a flagellar bundle behind the cell body. Thus, the elastic rod model is more accurate in describing the hook bending for motile bacteria.

### Measurements with the bead assay in a variable flow field

As shown in Fig. 2a, the cell body was adhered to the bottom surface of a microfluidic chamber, and a 0.5-*μm*-diameter bead was labeled to the truncated filament [29-31]. A steady fluid flow was supplied using a syringe pump with adjustable pump speed. As shown in Fig. 2b, the rotating bead exhibited a perfect circular trajectory in three dimensions (purple trajectory). The normal vector of the rotational plane N pointed to the tangential direction of the hook’s distal end. We recorded the projection of this trajectory on the image focal plane (XOY) via a high-speed CMOS camera.

**FIG. 2.**
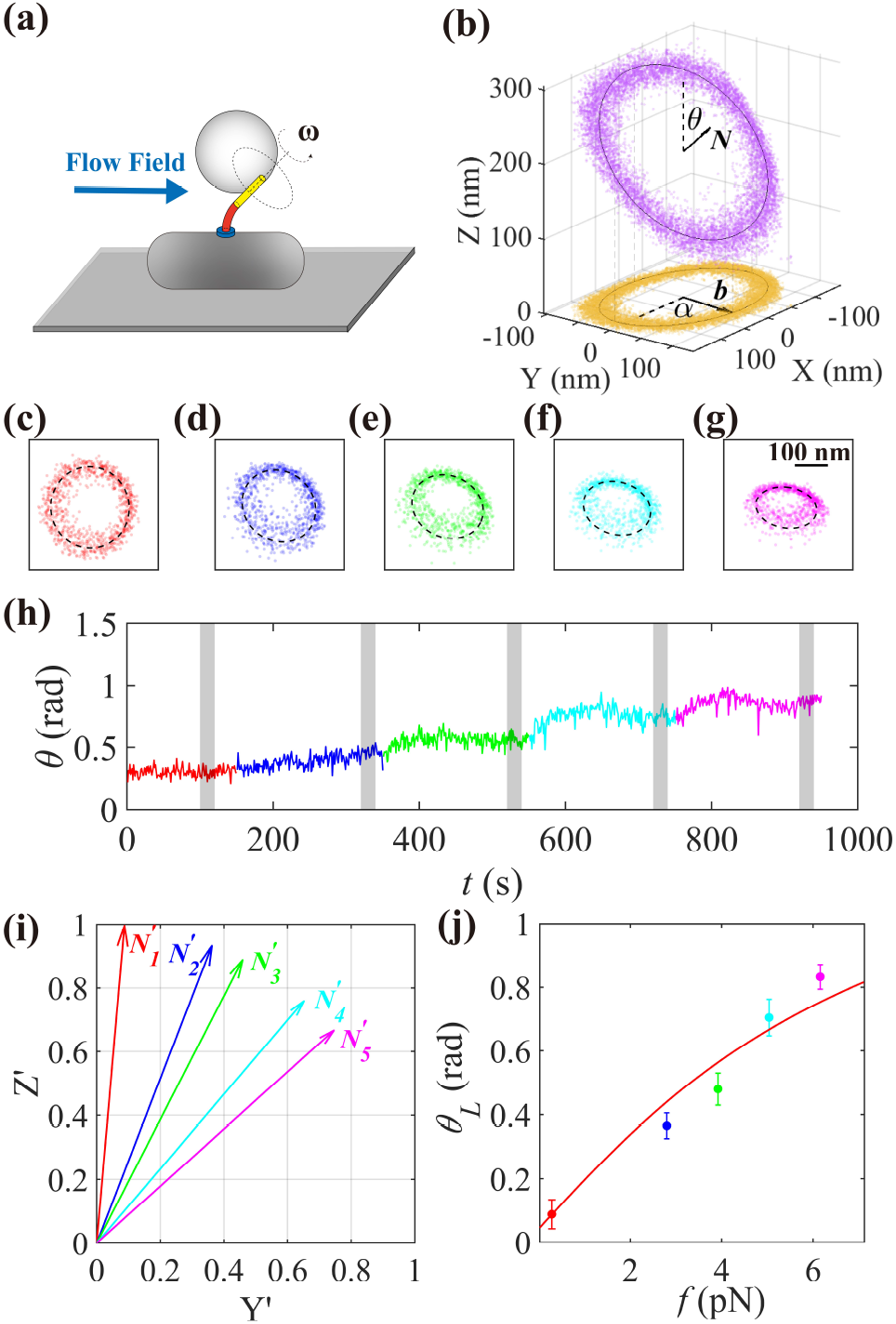
Typical measurement of hook bending stiffness in 10% Ficoll 400. **(a)** Schematic view of the experimental setup. A living bacterium was adhered to a microscope slide with a latex bead attached to the filament stub, and the medium was controlled by a syringe pump to generate a variable flow field. **(b)** Projection of 3D circular trajectories (purple) on the image focal plane XOY (yellow). **(c-g)** The 1-s-long projected trajectories of the rotating bead at pump speeds of 10 (red), 100 (blue), 140 (green), 180 (cyan) and 220 (magenta) *μL/min*. The black dashed lines denote the fitting results with an elliptic function. **(h)** The time trace of the tilted angle *θ* of the rotation plane with respect to the focal plane, calculated from the fitted ellipses. Five time windows were highlighted (gray regions) and considered as periods of steady state. Different colors represent the corresponding pump speeds, as denoted in (c-g). **(i)** The vectors 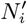 obtained by projecting the normal vectors of the bead rotating planes to the bending plane (Y’OZ’) for the selected time windows in (h). **(j)** The angle *θ*_*L*_ between 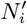 and the Z’-axis as a function of the flow force ***f***. The dots represent the experimental measurements, and the red solid line represents the fitted curve with the elastic rod model. The error bars denote SDs for the time periods.

To avoid the effect of motor switching, we chose two strains whose motors rotate exclusively in the CCW or CW direction. During each measurement, we recorded the rotation trajectories in steady flow fields with five different pump speeds. An example is shown in Fig. 2c-g, the bead trajectory gradually changes from a circle to an ellipse as the pump speed increases. By fitting these trajectories with an elliptic function, we can obtain the length of both the major (a) and the minor (b) axes, and the direction of the unit vector pointing along the short axis, (cos *α*, sin *α*). The ambiguity of the direction (0^*o*^ or 180^*o*^) was further resolved by following the motion trace of the ellipse center under different flow fields (Fig. S1). The tilted angle of the rotation plane with respect to the focal plane is *θ* = arccos(*b/a*). Typical evolution of *θ* over time for a motor with increasing external flow speeds is shown in Fig. 2h, exhibiting a stepwise increase as the hook bending increases with the flow speed. The hook is bent to equilibrium when *θ* enters a relatively stable plateau. We can now calculate the normal unit vector of the trajectory plane

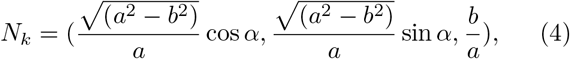

where *k* = 1, 2, 3, 4, or 5 denotes the index of the flow speeds. We selected five time-windows, each approximately 20 s, in the steady-state durations for the five different pump speeds (gray shaded areas in Fig. 2h). The hook bending plane could be obtained by fitting all *N*_*k*_ with a plane named Y’OZ’ (Fig. S2, see supplementary Note 2 for details), where the Y’-axis is the intersect of this plane with the XOY plane, and the Z’-axis is perpendicular to the Y’-axis and points to the positive direction of the Z-axis. More examples of hook bending traces are shown in Fig. S3. As all *N*_*k*_ do not perfectly lie in one plane, we projected *N*_*k*_ onto the Y’OZ’ plane to obtain 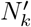 (Fig. 2i) and calculated the angle *θ*_*L*_ between 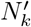 and the Z’-axis (Fig. 2j).

To quantify the fluid force applied on the bead, we should obtain the fluid flow speeds for different pump speeds. This was accomplished via a particle tracking velocimetry (PTV) or particle streak velocimetry (PSV) (see Supplemental Material for details), and the relations between the flow speeds v (*μm/s*) and pump speeds *V*_*p*_(*μL/min*) for motility medium with 0, 10% and 15% and 15% Ficoll 400 are shown in Fig. S4. By fitting with a linear function, we obtained *v*_0%*F icoll*_ = 1.55*V*_*p*_, *v*_10%*F icoll*_ = 1.47*V*_*p*_ and *v*_15%*F icoll*_ = 1.08*V*_*p*_. Thus, the force exerted by the flow field on the bead can be computed as

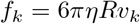

where *η* is the media viscosity, *R* denotes the bead radius, and *v*_*k*_ represents the flow speed of the solution around the bead. An example of the relation between *θ*_*L*_ and f measured for a motor is plotted in Fig. 2j. We evaluated the length of filament stub *L*_*stub*_ for each motor by combining the movements of the center of the trajectory with the changes in *N*_*k*_ as the flow field changed (see Supplemental Material for details). Using equation (3) with the values of *L*_*stub*_, *f* and *θ*_*L*_ extracted above, and with *L*_*hook*_ = 55*nm*, the parameters *EI* and *θ*0 can be obtained by fitting the data in Fig. 2j (see supplementary Note 3 for details). We obtained *EI* = 4.73×10^−^25*N* · *m*2 and *θ*0 = 0.042 *rad* for this motor from the fitting.

### Comparison of hook bending stiffness for CW and CCW-rotating motors

To compare the bending stiffness of the hook in the CW and CCW-rotating states, we performed the measurements for both the CCW and CW strains. Considering the load dependence of the hook bending stiffness, three typical loads were selected by adding 0%, 10% or 15% (w/v) Ficoll 400 to the motility medium. We measured a total of 135 motors.

The results are shown in Fig. 3a. The order of magnitude of the bending stiffness we measured is ∼ 10^−25^*N* · *m*^2^ which is compatible with previous analyses[9,26]. Specifically, the mean values of the bending stiffness of the hooks under different conditions were measured to be:

**FIG. 3.**
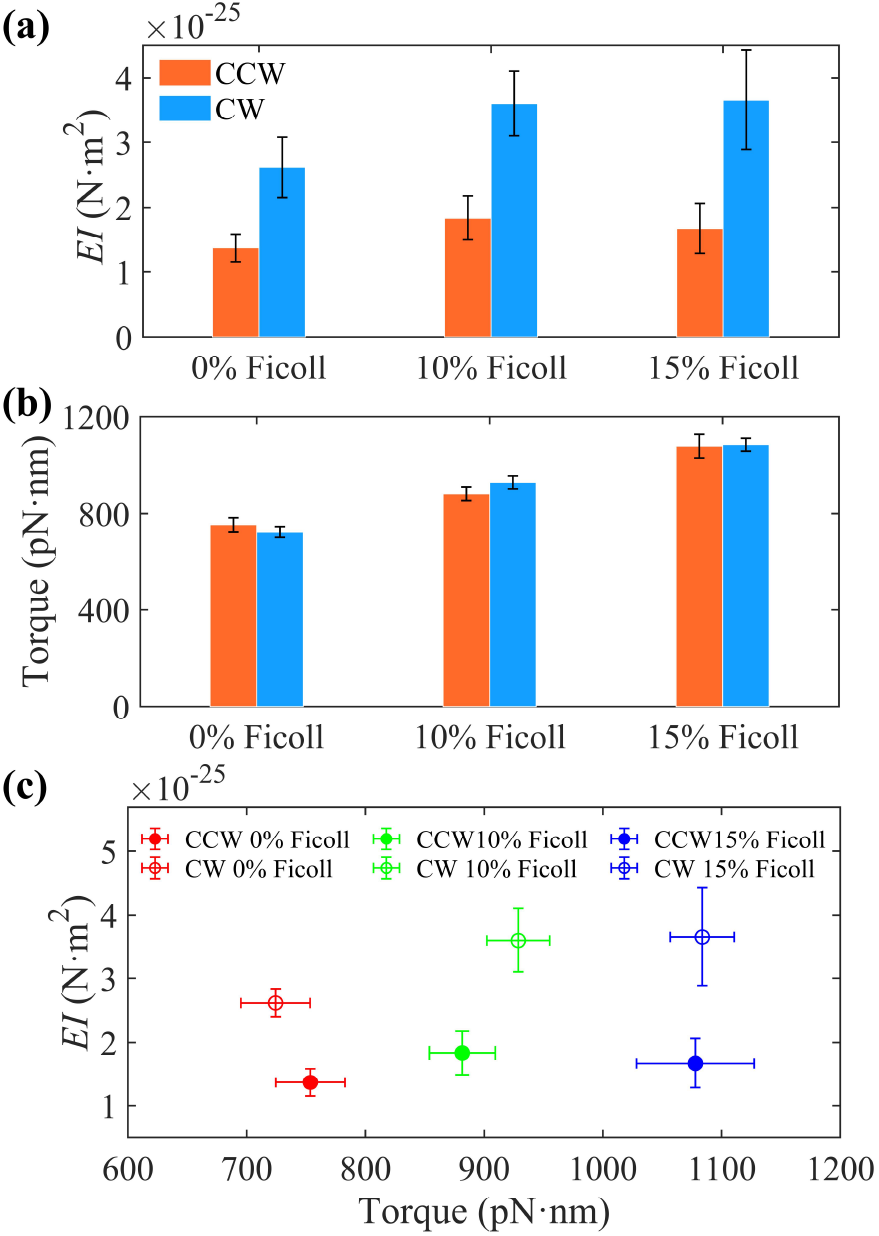
**(a)** The hook bending stiffness for the CCW- and CW-rotating strains in 0%, 10% and 15% Ficoll 400 solutions. The error bar denotes the standard error of the mean (SEM). The motor torque for CCW- and CW-rotating strains in 0%, 10% and 15% Ficoll 400 solutions. **(c)** The hook bending stiffness versus the motor torque under different conditions. The error bar denotes SEM.

1. In 0% Ficoll 400,

*EI*_*CCW*_ = (1.37 ± 0.22) × 10^−25^*N* · *m*^2^,

*EI*_*CW*_ = (2.62 ± 0.46) × 10^−25^*N* · *m*^2^.

1. 3.In 10% Ficoll 400,

*EI*_*CCW*_ = (1.84 ± 0.34) × 10^−25^*N* · *m*^2^,

*EI*_*CW*_ = (3.60 ± 0.50) × 10^−25^*N* · *m*^2^.

1. In 15% Ficoll 400,

*EI*_*CCW*_ = (1.67 ± 0.39) × 10^−25^*N* · *m*^2^,

*EI*_*CW*_ = (3.75 ± 0.75) × 10^−25^*N* · *m*^2^.

We note that the mean bending stiffness of hooks in the CW strain is 1.9 ∼ 2.2 times larger than the CCW strain under the same load condition.

We calculated the torque for each motor using the drag coefficients of the bead *f*_*b*_ = 8*πη*^3^ + 6*πηRs*^2^ [30], where *η* is the media viscosity measured by a viscometer [32], *R* = 0.25*μm* is the bead radius, and s denotes the distance between the bead center and the rotational axis. We also estimated the contribution of the filament stub to the drag coefficients using a method similar to that used previously [33] (see Supplemental Material for details). As shown in Fig. 3b and 3c, the torques are roughly the same for the two rotational directions under the same load conditions; thus, the difference in the hook bending stiffness for the two rotational directions is not caused by the difference in loads.

## Discussion

Here, we investigated the hook bending stiffness of *E. coli* when the motors rotated in the CCW or CW direction under different loads by using bead assays with variable flow fields. The ranges of the bending stiffness *EI* we measured are 1.37 ×10^−25^ ∼ 1.84 ×10^−25^*N* · *m*^2^ and 2.62 ×10^−25^ ∼3.75 ×10^−25^*N* · *m*^2^ for CCW and CW rotations, respectively.

These results demonstrated that the bending stiffness of the hook when the motor rotates CCW is twice smaller than that when the motor rotates CW. The differential bending stiffness of the hook can effectively promote the bundling of the filaments behind the cell for smooth swimming when the motors rotate CCW, and meanwhile promote unbundling of the filaments when the motors rotate CW.

The structure of the bacterial hook is considered to be a hollow cylinder as measured by cryoelectron microscopies [3,14,34], and the area moment of inertia I can be calculated from the cross section of the hollow cylinder as 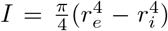, where *r*_*e*_ = 9 *nm* and *r*_*i*_ = 1.5 *nm* are the external and internal radii, respectively. Using our measured values of *EI*, the Young’s modulus E is extracted to be ∼ 10^7^*dyn/cm*^2^, consistent with the range of theoretical predictions [1].

Our study showed that the effect of twist direction on the bending stiffness is asymmetric, and the bending stiffness under CW twist is higher than that under CCW twist for the same torsional stress. This may be understood qualitatively by the polymorphic transformation of the hook triggered by switching of the flagellar motor. The hook as well as the filament can transform among a series of helical forms when the external conditions of temperature, pH and ionic strength change [35]. Considering that the flagellar filament with structure similar to the hook changes its supercoil structure under the induction of flagellar motor torque [36-39], it is reasonable to infer that the hook can also change its polymorphic forms under motor torque. Furuta et al. investigated the hook structure by molecular dynamics simulation, and observed two extreme cases, the left-handed and right-handed coils, which make maximal use of the gaps between subunits [40]. Interestingly, they found that the right-handed coil subunits are much more tightly packed compared to the left-handed coil; that is, the left-handed coil packing reserve more gaps for further compression [40]. The microscopic differences between the left- and right-handed coils of the hook may lead to different mechanical properties. The differential bending stiffness of the hook we measured here may correspond to different polymorphic forms of the hook under CW and CCW rotations. Further studies are needed to elucidate the detailed underlying molecular mechanism of the differential bending stiffness.

This work was supported by National Natural Science Foundation of China Grants (11925406, 11872358, and 12090053), a grant from the Ministry of Science and Technology of China (2019YFA0709303), and a grant from the Natural Science Foundation of Anhui Province (2008085QA31).

