## Supplemental Material for "Differential bending stiffness of the bacterial flagellar hook under counterclockwise and clockwise rotations"

### Materials and methods

**Strains and plasmids.** Strains JY27 (*ΔfliC cheY*) and XL2(*ΔfliC cheY cheZ cheB*) are derived from the *E. coli* K12 strain RP437. The plasmid pKAF131 constitutively expresses the sticky *fliC* allele, which was cloned into a pACYC184 ($Cam^R$) vector under the native promoter of *fliC* [1]. The plasmid pWB5 ($Amp^R$) expresses wild-type CheY under an IPTG-inducible promoter [2,3]. For measurements of the bending stiffness for the CCW motor, JY27 carrying pKAF131 was used. For measurements of bending stiffness for the CW motor, XL2 transformed with pKAF131 and pWB5 was used. The CW bias distributions for both strains are shown in Fig. S5.

**Cell culture.** Cells of JY27 transformed with pKAF131 were grown at 33°C in T-broth with the appropriate antibiotics (25 μg/mL chloramphenicol). Cells of XL2 transformed with pWB5 and pKAF131 were grown in T-broth with the appropriate antibiotics (25 μg/mL chloramphenicol and 100 μg/mL ampicillin) and the appropriate inducers (200 μM IPTG for CheY expression). All of them were grown to an $OD_{600}$ between 0.45 and 0.50 and were harvested by centrifugation at 4000×*g* for 2 min twice with motility medium (10 mM potassium phosphate, 0.1 mM ethylenediaminetetraacetic acid and 10 mM lactic acid (pH 7.0)) and resuspended in this medium. Cells were used immediately or stored at 4°C for up to 3.5 h.

**Bead assay.** Cells were sheared to truncate flagella by passing 1 mL of the washed-cell suspension 120 times between two syringes equipped with 23-gauge needles and connected by a 16-cm length of polyethylene tubing (0.58 mm i.d., 0.965 mm o.d., no. 427411; Becton Dickinson, Franklin Lakes, NJ). Coverslips were coated with poly-L-lysine (0.01%, catalog no. P4707; Sigma). Each glass slide we used was drilled with two 1 mm diameter holes that were glued with structural plastic adhesive (DP8005 off-white; Scotch-Weld, 3M, Saint Paul, MN) to two 3.5-cm lengths of polyethylene tubing (0.58 mm i.d., 0.965 mm o.d., no. 427411; Becton Dickinson, Franklin Lakes, NJ). A chamber was formed with a circular double-sided tape sealed with vacuum grease between the coverslip and the glass slide, including the two holes and polyethylene tubing described above. To measure the motor rotation in the CCW or CW direction, 80 μL cells were added to the chambers from one of the polyethylene tubes, allowed to stand for 5 min, and rinsed with 200 μL motility medium to wash away free cells. To measure the motor rotation under different loads, 0.5-μm-diameter polystyrene latex beads (Polysciences, Warrington, PA) were attached to the sheared filament stubs, incubated for 4 min, and rinsed with 200 μL motility medium. The polystyrene beads were observed using a Nikon Ti-E inverted phase-contrast microscope. The chamber was connected to a vessel filled with medium solution and a syringe pump to control the flow field.

The medium solutions we selected to generate the flow field were 0%, 10% and 15% (w/v) Ficoll 400 in motility medium (the viscosity was measured to be 1.0 cP, 3.8 cP and 7.0 cP, respectively). The pumping speed settings are listed in table S1. Phase-contrast images were recorded at frame rates of 498 ~ 1980 frames per second (fps) using a scientific complementary metal-oxide semiconductor (CMOS) camera (C11440; Hamamatsu, Hamamatsu, Japan). Data analysis was carried out using custom scripts in MATLAB (The MathWorks, Natick, MA). A total of 135 motors were measured. Specifically, for the CCW stain, 19 motors for cells in 0% Ficoll, 22 motors for cells in 10% Ficoll, and 27 motors for cells in 15% Ficoll were measured. For the CW strain, 19 motors for cells in 0% Ficoll, 20 motors for cells in 10% Ficoll, and 28 motors for cells in 15% Ficoll were measured.

**Particle tracking velocimetry and particle streak velocimetry.** Flow velocities in the chamber near the coverslip surface, where the cells were stuck, were measured using particle tracking velocimetry (PTV) [4,5] or particle streak velocimetry (PSV) [6-8]. Using 1-μm-diameter latex beads as tracer particles, the original bead solution was diluted to 0.27% and then was mixed with 0%, 10% or 15% Ficoll 400 solution in a 1:6 volume ratio, yielding the medium solution labeled by beads. We used the same flow device as the bead assay with the medium solution labeled by beads, and the pump speeds were set to be the same as that used in the bead assay. The tracer particles were observed by a Nikon Ti-E inverted phase-contrast microscope at a magnification of 40× and recorded with a scientific CMOS camera (C11440; Hamamatsu, Hamamatsu, Japan). For the PTV method, we adjusted the frame rate (30-500 fps) to capture the motion trace of particles in the plane of rotating beads, and the velocity of the flow field was obtained by calculating the average value of the displacements of the tracer particles between two adjacent frames multiplied by framerate (as shown in Fig. S6). When the translational speeds of the tracer particles were relatively fast, we used PSV to widen the range of interest to capture more tracer particles. For the PSV method, we adjusted the exposure time to generate particle streaks. As shown in Fig. S7, the velocity of the flow field was obtained by calculating the average value of the trajectory lengths of the tracer particles moving in the exposure time divided by the exposure time interval. Data analysis was carried out using custom scripts in MATLAB (The MathWorks, Natick, MA).

**Measurement of the length of the filament stub.** Assuming that the movement of the center of the bead trajectory is mainly caused by the bending of the hook, the length of the filament stub can be estimated by calculating the projection of the change of the unit normal vector on the XOY plane (Fig. 2). After applying the drift correction [9] to the trajectory (see *Supplementary Note 4*), we calculate the change in the unit normal vector from the trajectories of the bead in different flow fields, $\Delta \boldsymbol{N_j} = \boldsymbol{N_{j+1}} - \boldsymbol{N_j}$ $(j = 1,2,3,4)$, and the displacement $\Delta \boldsymbol{x_j}$ of the center coordinates $C_j$ of the trajectory, $\Delta \boldsymbol{x_j} = C_{j+1} - C_j$. The length of the filament stub $l_j$ can be computed as $l_j = |\Delta \boldsymbol{x_j}| / |\Delta \boldsymbol{N_j} - (\Delta \boldsymbol{N_j} \cdot \boldsymbol{e_z}) \boldsymbol{e_z}|$, where $\boldsymbol{e_z}$ is the unit normal vector of the XOY plane. We calculated the average value of $l_j$ for each motor as the length of the flagella stub. The length distribution of the filament stubs for all motors is shown in Fig. S8.

**Measurement of the motor torque.** The rotational frictional drag coefficients are $f = f_b + f_f$, where $f_b$ and $f_f$ are the drag coefficients of the beads and the filament stubs, respectively. For a sphere of radius $a$ rotating about an axis offset by a distance $l$ from its center in a medium of viscosity $\eta$, the drag coefficients of the bead $f_b = 8\pi\eta a^3 + 6\pi\eta a l^2 = g_b\eta$, where $g_b$ is a geometry factor of the bead. Similarly, $f_f = g_f\eta$, where $g_f$ is a geometry factor of the filament stubs and is considered to be a constant. We obtain $g_f$ by equating the plateau torques measured by attaching 1-μm-diameter bead to the filament stub in motility medium and by attaching 0.5-μm-diameter bead to the filament stub in 15% Ficoll 400 using a similar method as before , and the value of $g_f$ is estimated to be $(4.34\pm0.97)\times10^7$ $nm^3$ (mean±SD, see *Supplementary Note 5*). The torque exerted on the bead by viscous drag is $f\Omega$, where $\Omega$ is the angular velocity of the bead (2π times its rotation speed in Hz).

**Supplementary Note 1: Detailed derivation of the equation of bending stiffness of the bacterial hook in bead assay.** In classical elastic theory [10], bending and twisting of a rod with a circular cross-section are independent. Therefore, we consider that the hook undergoes pure bending. We establish a fixed reference coordinate system (O-XYZ) and an arc coordinate $l$ along the rod in the centerline, and the position of any point on this line is determined by the radius vector $\boldsymbol{r}$ in (O-XYZ) or the arc coordinate $l$.

For an infinitesimal element bounded by two adjoining cross-sections of the rod in equilibrium and the distance between two cross-sections is infinitesimal $dl$, the equations of resultant internal stress $\boldsymbol{F}$ and the moment of internal stress $\boldsymbol{M}$ in equilibrium for rod bending are as follows:

$$\frac{d\boldsymbol{F}}{dl} = -\boldsymbol{K}, \tag{1}$$

$$\frac{d\boldsymbol{M}}{dl} = \boldsymbol{F} \times \boldsymbol{t}, \tag{2}$$

where $d\boldsymbol{F}$ and $d\boldsymbol{M}$ represent the change of $\boldsymbol{F}$ and $\boldsymbol{M}$ between the upper and lower cross-section of the infinitesimal element, $\boldsymbol{K}$ is the external force on the rod per unit length, and $\boldsymbol{t}$ is the tangent vector of any point on the rod ($\boldsymbol{t} = d\boldsymbol{r}/dl$). Since the radius of the rod section (~10 nm) is much smaller than that of the bead (~250 nm), the flow force on the rod is negligible ($\boldsymbol{K} = 0$). For a rod with a circular cross-section in pure bending, we can derive an expression for $\boldsymbol{M}$:

$$\boldsymbol{M} = IE\boldsymbol{t} \times \frac{d\boldsymbol{t}}{dl} = IE\frac{d\boldsymbol{r}}{dl} \times \frac{d^2\boldsymbol{r}}{dl^2} \tag{3}$$

where $E$ is the Young's modulus and $I = I_1 = I_2$ are the principal moments of inertia for a rod with a circular cross-section. Substituting eq. (3) in eq. (2), we obtain

$$IE\frac{d\boldsymbol{r}}{dl} \times \frac{d^3\boldsymbol{r}}{dl^3} = \boldsymbol{F} \times \frac{d\boldsymbol{r}}{dl} \tag{4}$$

We create a right-handed Cartesian coordinate system O-X′Y′Z′, where the Y′-axis is parallel to the flow field force, the Z′-axis is in the bending plane and perpendicular to the Y′ axis pointing to the positive direction of the Z-axis, and the X′-axis is perpendicular to the Y′OZ′ plane. The basis vectors of O-X′Y′Z′ are $\boldsymbol{e}_{x'}$ ,$\boldsymbol{e}_{y'}$, and $\boldsymbol{e}_{z'}$. In this coordinate system, we have $\boldsymbol{F} = (0, f, 0)$ and $\boldsymbol{t} = \frac{dr}{dl} = (0, sin\theta, cos\theta)$, where $f$ is the magnitude of the flow force and $\theta$ is the angle between the tangent line of any point on the rod and the Z′-axis. The second and third derivatives of $\boldsymbol{r}$ are

$$\frac{d^2\boldsymbol{r}}{dl^2} = \left(0, cos\theta\frac{d\theta}{dl}, -sin\theta\frac{d\theta}{dl}\right) \tag{5}$$

$$\frac{d^3\boldsymbol{r}}{dl^3} = \left(0, -sin\theta\left(\frac{d\theta}{dl}\right)^2 + cos\theta\frac{d^2\theta}{dl^2}, -cos\theta\left(\frac{d\theta}{dl}\right)^2 - sin\theta\frac{d^2\theta}{dl^2}\right) \tag{6}$$

Combining eqs. (3), (4), (5) and (6), together with the expressions of $\boldsymbol{F}$ and $\boldsymbol{t}$, we can obtain the equation of pure bending of a rod with a circular section,

$$IE\frac{d^2\theta}{dl^2} + fcos\theta = 0 \tag{7}$$

$$M = -IE\frac{d\theta}{dl} \tag{8}$$

Taking the first integral of eq. (7) gives $\frac{1}{2}IE\left(\frac{d\theta}{dl}\right)^2 + fsin\theta = c_1$, and further integration leads to

$$l = \pm\sqrt{\frac{IE}{2}}\int\frac{d\theta}{\sqrt{c_1 - fsin\theta}} + c_2 \tag{9}$$

The boundary conditions are $\theta = \theta_0$ for $l = 0$ at the clamped end, and at the connecting point of the hook and the filament stub $\theta(L_{hook}) = \theta_L$, $\left.\frac{d\theta}{dl}\right|_{l=L_{hook}} = -\frac{fL_{stub}cos\theta_L}{IE}$ since $M = fL_{stub}cos\theta_L$ for $l = L_{hook}$, where $L_{hook}$ is the length of the hook, and $L_{stub}$ is the length between the point where the bead sticks and the connecting point of the hook and the filament stub. Then, we have

$$L_{hook} = \sqrt{\frac{EI}{2f}} \int_{\theta_0}^{\theta_L} \frac{d\theta}{\sqrt{\frac{fL_{stub}^{\ 2}\cos^2\theta_L}{2EI} + \sin\theta_L - sin\theta}}. \qquad (10)$$

**Supplementary Note 2: Computation of the normal vector of the bead rotation plane.** The measured trajectories of the beads are scarcely perfectly circular in bead assay, because we only recorded the projection of the 3-d trajectory of the rotating bead on the image focal plane (XOY). Assuming that the trajectory is circular before projection to the focal plane [11,12], we calculate the normal vectors of the beads' rotational plane using the parameters of the fitted ellipse. In Fig. S9 (a), we show an elliptical trajectory with the fitted ellipse.

The length of the major axis $a$ and minor axis $b$ of the ellipse and the inclination angle of the short axis $\alpha$ can be obtained from the parameters of the fitted ellipse. Since we assume a circle before the projection, the projection on the XOY plane of the normal vector is parallel to the minor axis, and the cosine of the angle $\theta$ between the normal vector and the Z-axis is $a/b$ (Fig. S9(b-d)). To eliminate degeneracy, we performed a normal vector calculation for each second of the bead traces and chose the *x* and *y* components to ensure that the normal vector change for the consecutive two seconds was not more than $\frac{\pi}{2}$, and the *z* component was specified to be positive (that is, $0 \leq \theta \leq \frac{\pi}{2}$). After normalization, the normal vector ***N*** can be calculated as:

$$\boldsymbol{N} = (sin\theta cos\alpha, sin\theta sin\alpha, cos\theta) \qquad (11)$$

**Supplementary Note 3: Parameter fitting of $\boldsymbol{EI}$ and $\boldsymbol{\theta_0}$ for each motor using a nonlinear solver.** Using the obtained parameters $\theta_k{}'$ and $f_k$ $(k = 1,2,3,4,5)$ to substitute $\theta_L$ and $f$ in eq. (10), respectively, as well as $L_{hook}$ and $L_{stub}$, five calculated results of the hook length were obtained under different flow fields, and then the following function was obtained:

$$MSE(\theta_0, EI) = \frac{1}{5}\sum_{k=1}^{5}\left(\sqrt{\frac{EI}{2f_k}}\int_{\theta_0}^{\theta'_k}\frac{d\theta}{\sqrt{\frac{f_k L_{stub}{}^2 \cos^2\theta'_k}{EI} + \sin\theta'_k - \sin\theta}} - L_{hook}\right)^2 \quad (12)$$

which is the **m**ean **s**quared **e**rror (MSE) between the calculated values and measured value of the length of the hook. For each motor, the optimal values for $\theta_0$ and $EI$ are found by unconstrained nonlinear minimization (with the MATLAB function `fminsearch`) of the value of $MSE(\theta_0, EI)$.

**Supplementary Note 4: Drift correction for the sample stage.** We corrected the lateral drift of the sample stage by tracking the center of the beads' rotational trajectories in the steady states. We selected five time windows as steady states in the time periods of different flow rates when $\theta$, which is the tilted angles of the rotation planes with respect to the focus plane, is relatively stable (Fig. S1a). For simplicity, we considered only linear drift with constant drift velocity components $V_{x,k}$ and $V_{y,k}$ in each steady period $k$, which were calculated from the change of the trajectory centers per unit time for each steady period (Fig. S1b). The drift velocities in the time periods between two steady states $k$ and $(k+1)$ are also considered to be constant with drift velocity components $(V_{x,k} + V_{x,k+1})/2$ and $(V_{y,k} + V_{y,k+1})/2$. The drift displacement of the sample stage at each time point can then be calculated by integration/summation using the drift velocity components, and the drift correction can be applied by subtracting the drift displacement (Fig. S1c-d).

**Supplementary Note 5: Estimation of the geometry factor $\boldsymbol{g_f}$ of the filament stub.** Assuming the geometric factor $g_f = f_f/\eta$ for the drag coefficient of the filament stub to be constant, we obtain $g_f$ by equating the plateau motor torques measured at low speed by 1-μm-diameter beads in motility medium and by 0.5-$\mu$m-diameter beads in 15% Ficoll 400 using a method as before [13]:

$$g_f = \frac{g_{b2}\eta_2\omega_2 - g_{b1}\eta_1\omega_1}{\eta_1\omega_1 - \eta_2\omega_2} \tag{13}$$

where $g_{b1}$ and $g_{b2}$ are geometric factors of 1- and 0.5-μm beads, $\eta_1$ $(= 1.0 \times 10^3\ Pa \cdot s)$ and $\eta_2$ $(= 7.0 \times 10^3\ Pa \cdot s)$ are viscosities of motility medium and 15% Ficoll 400, and $\omega_1$ and $\omega_2$ are speeds measured with 1-μm-diameter beads in motility medium and 0.5-μm-diameter beads in 15% Ficoll 400, respectively. The average value of $g_f$ with SD is $g_f = (4.34 \pm 0.97) \times 10^7\ nm^3$. For comparison, the average values for $g_{b1}$ and $g_{b2}$ are $(3.50 \pm 0.06) \times 10^9\ nm^3$ and $(4.64 \pm 0.12) \times 10^8\ nm^3$, respectively.

**Supplementary tables**

**Tab. S1.** Flow rate settings for the microfluidic device

| **Strain** | **Ficoll 400 (w/v)** | **Pumping speed (μL/min)** | | | | |
|---|---|---|---|---|---|---|
| | | 0~150s | 150~350s | 350~550s | 550~750s | 750~950s |
| **CCW** | 0% | 10 | 400 | 560 | 720 | 880 |
| **CCW** | 10% | 10 | 100 | 140 | 180 | 220 |
| **CCW** | 15% | 10 | 100 | 140 | 180 | 220 |
| **CW** | 0% | 10 | 600 | 840 | 1080 | 1320 |
| **CW** | 10% | 10 | 200 | 280 | 360 | 440 |
| **CW** | 15% | 10 | 100 | 140 | 180 | 220 |

**Supplementary figures**

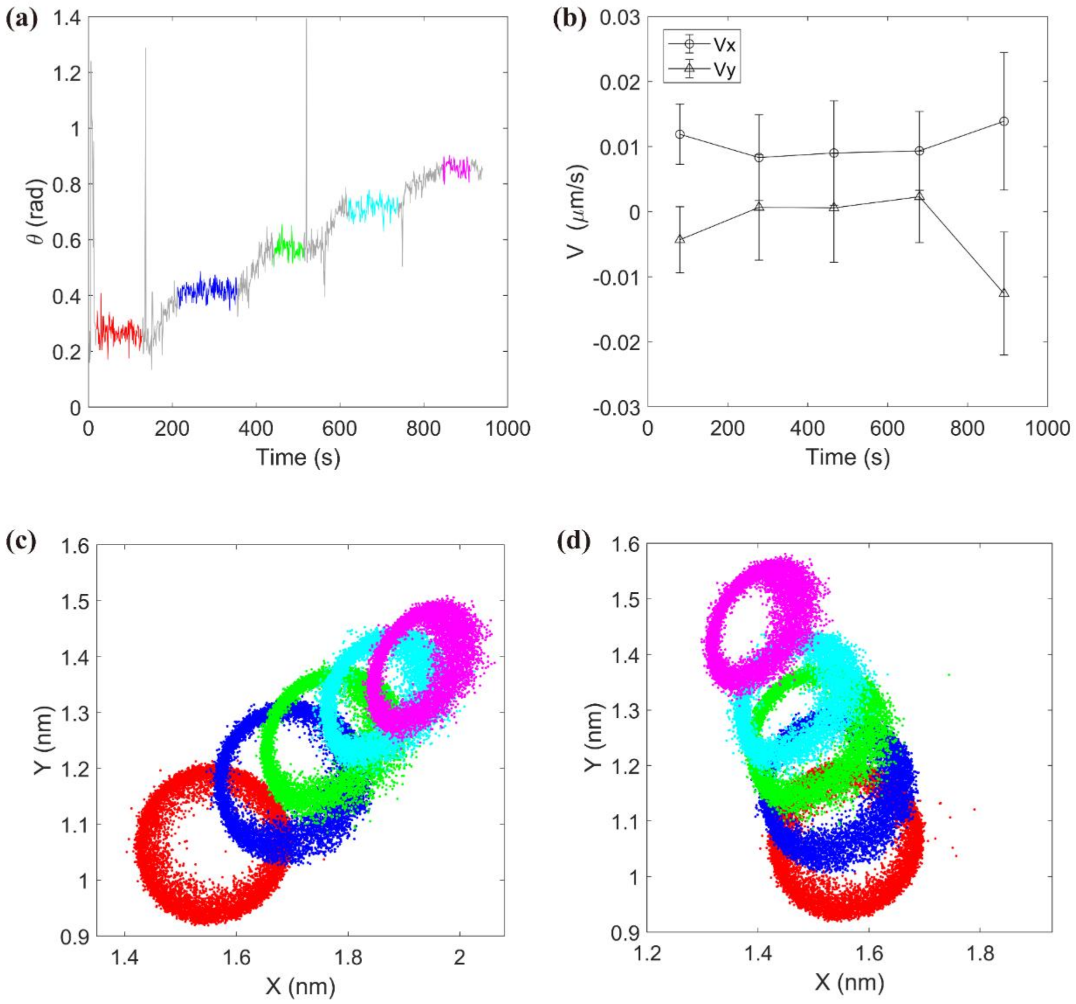


**Fig. S1.** An example of drift correction. **(a)** The change of tilted angles $\theta(t)$ of the rotational planes with respect to the focal plane over time. Five time windows under different flow fields are highlighted (red, blue, green, cyan and magenta represent the corresponding flow fields as shown in Fig. 2h). **(b)** The drift velocities along the X-axis and Y-axis calculated for the highlighted time windows in (a). **(c)** The trajectories of the bead before correction. **(d)** The trajectories of the bead after correction. The trajectories of the bead in (c) and (d) are taken from the highlighted time windows in (a) indicated by the corresponding colors.

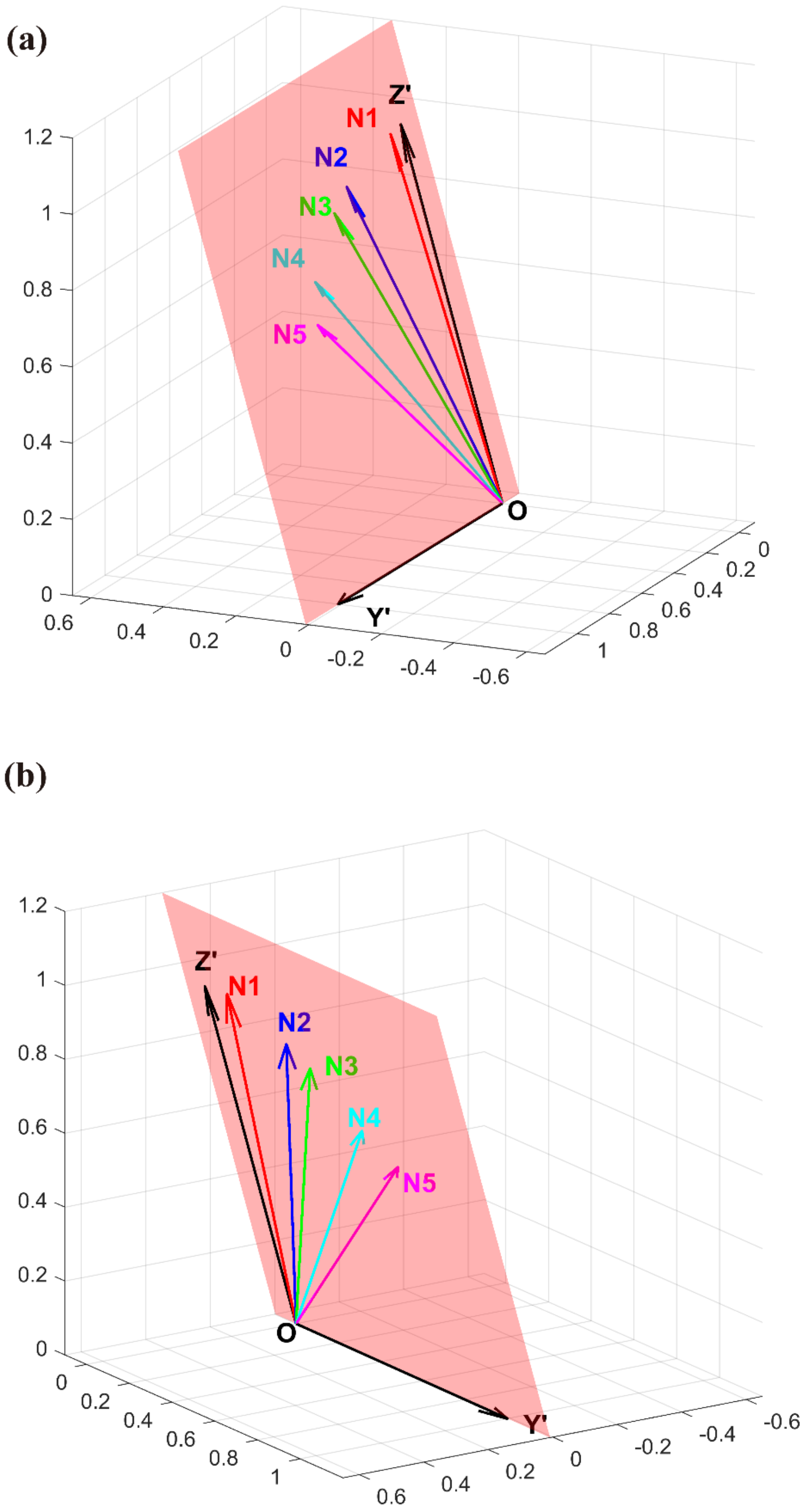


**Fig. S2.** The normal vectors $\boldsymbol{N}_k$ of the bead rotating planes and the fitted bending plane (Y′OZ′, red shaded region) for the example listed in Fig. 2, where (a) and (b) are viewed from different perspectives.

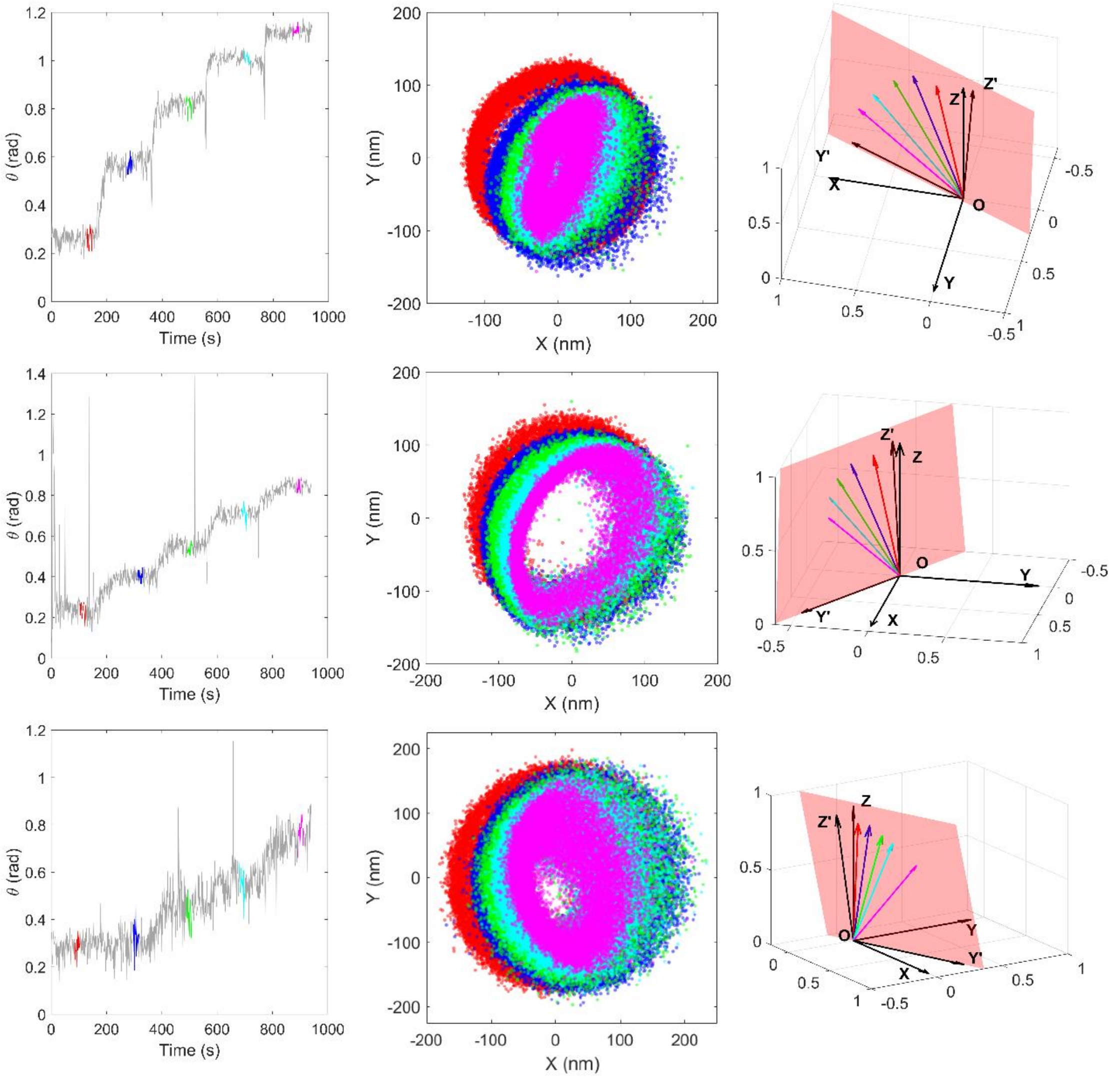


**Fig. S3.** Examples of trajectory processing of the bead assay. Each row corresponds to a different bead under different conditions, which are 0% Ficoll, 10% Ficoll and 15% Ficoll from top to bottom. The left column shows the tilted angles $\theta(t)$ of the rotation planes with respect to the focal plane, which were calculated from the fitted ellipse of every 1-s trace in the whole trajectory. Five time windows, each 20 s long, were highlighted (red, blue, green, cyan and magenta), which were considered as periods of time in steady states. The middle column shows the bead trajectories in the highlighted time windows with the corresponding colors. The right column shows the normal vectors obtained from the bead trajectories with the corresponding colors and the bending plane (red shaded region).

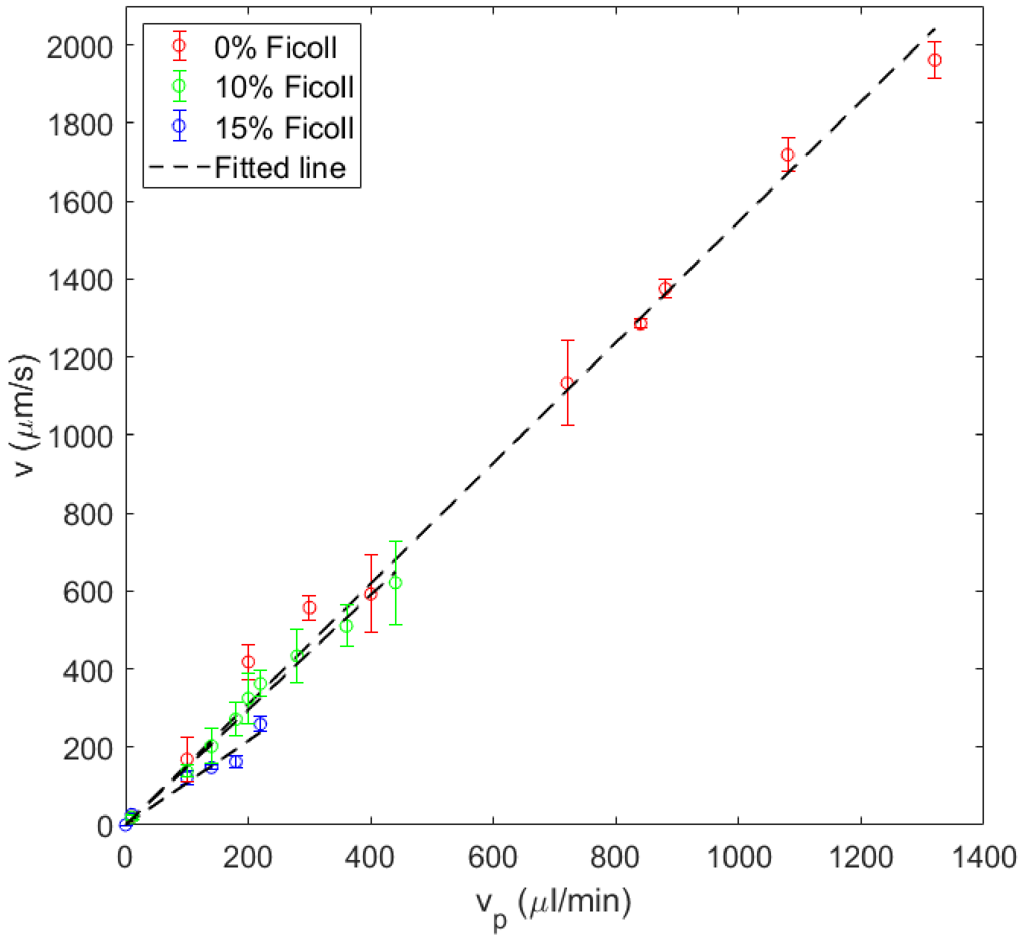


**Fig. S4.** The average speed of the flow field near the focal plane under different pump speeds, where different colors represent different concentrations of Ficoll 400 solutions and the dashed lines represent linear fits.

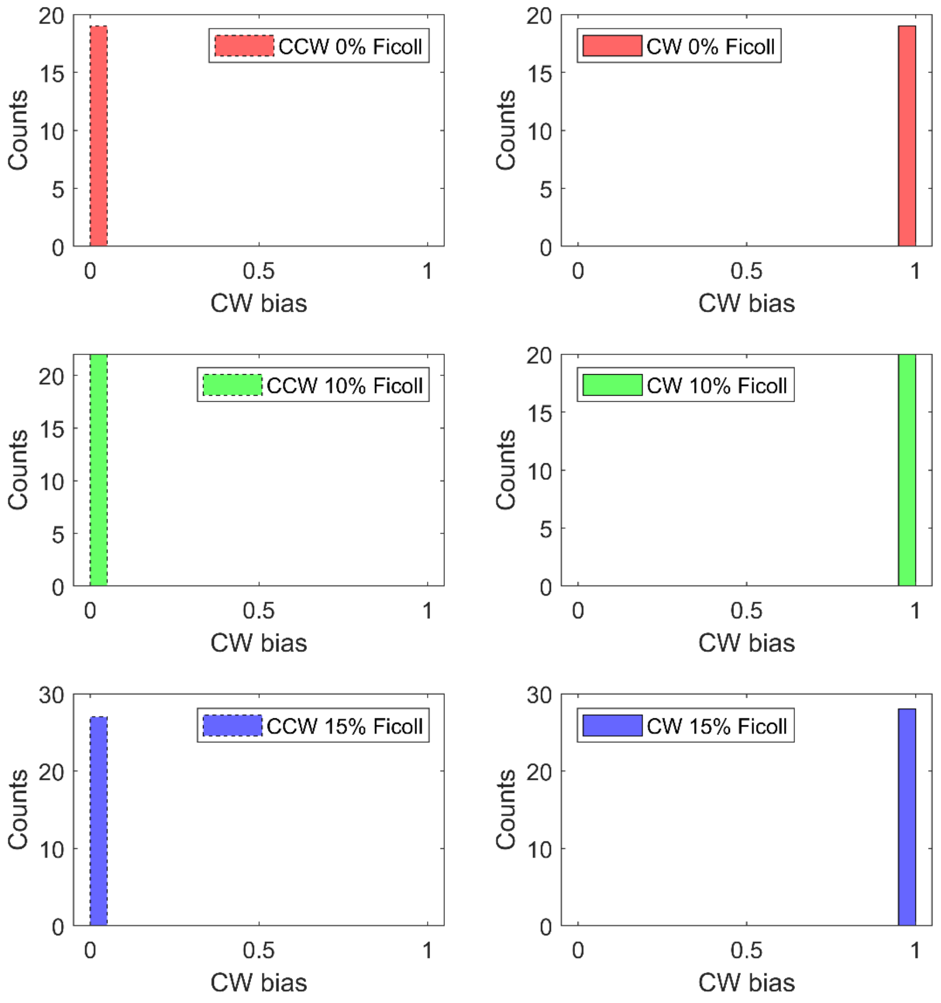


**Fig. S5.** The CW bias distribution of CCW- and CW-rotating strains in different concentrations of Ficoll 400.

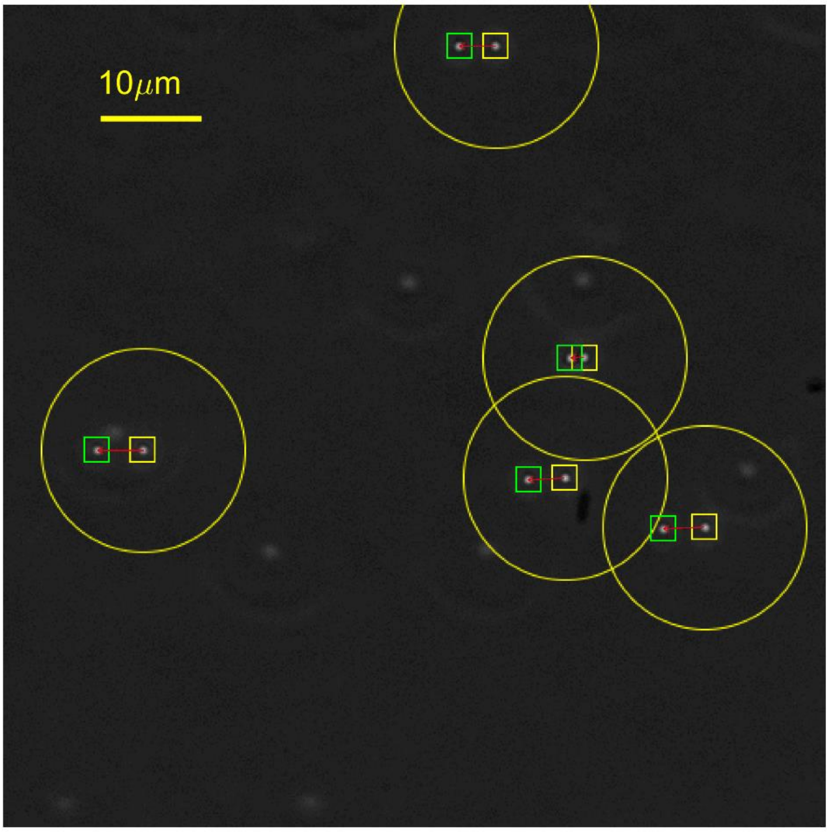


**Fig. S6.** Image processing of PTV applied to the sequence of two adjacent frames (from $i$ to $i+1$). Detected particles in frames ($i$) and ($i+1$) are marked with yellow and green squares, respectively. Each particle in frame ($i$) has a search radius of 10 μm (yellow circle) to find a unique matching particle in frame ($i+1$), and the red arrows denote the velocity vectors of the particles in frame ($i$).

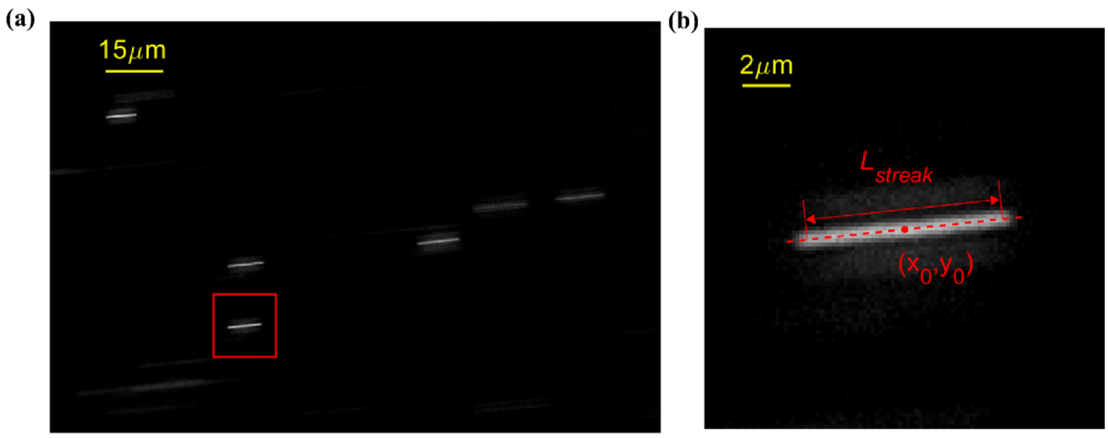


**Fig. S7.** Example of image processing in particle streak velocimetry. **(a)** A streak image from PSV. **(b)** The processing implementation of PSV, the length of the streak $l_{streak}$ and the centroid $(x_0, y_0)$ were obtained by fitting of the gray scale distribution.

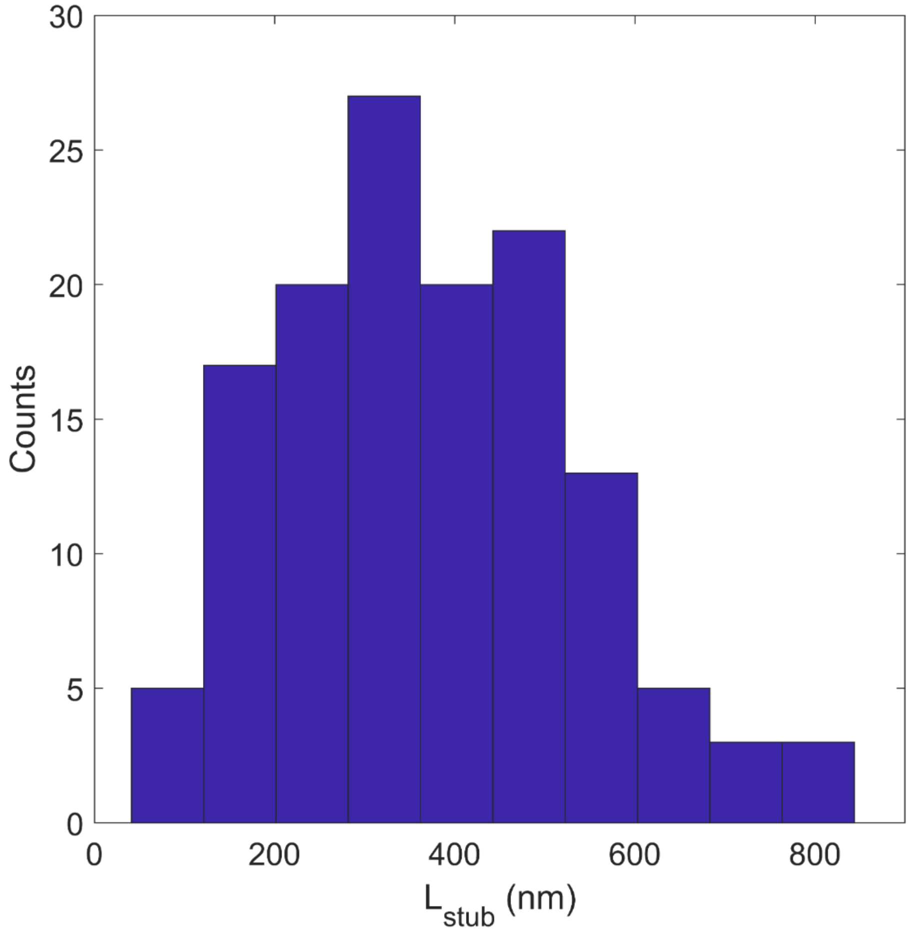


**Fig. S8.** The length distribution of the filament stubs for all motors.

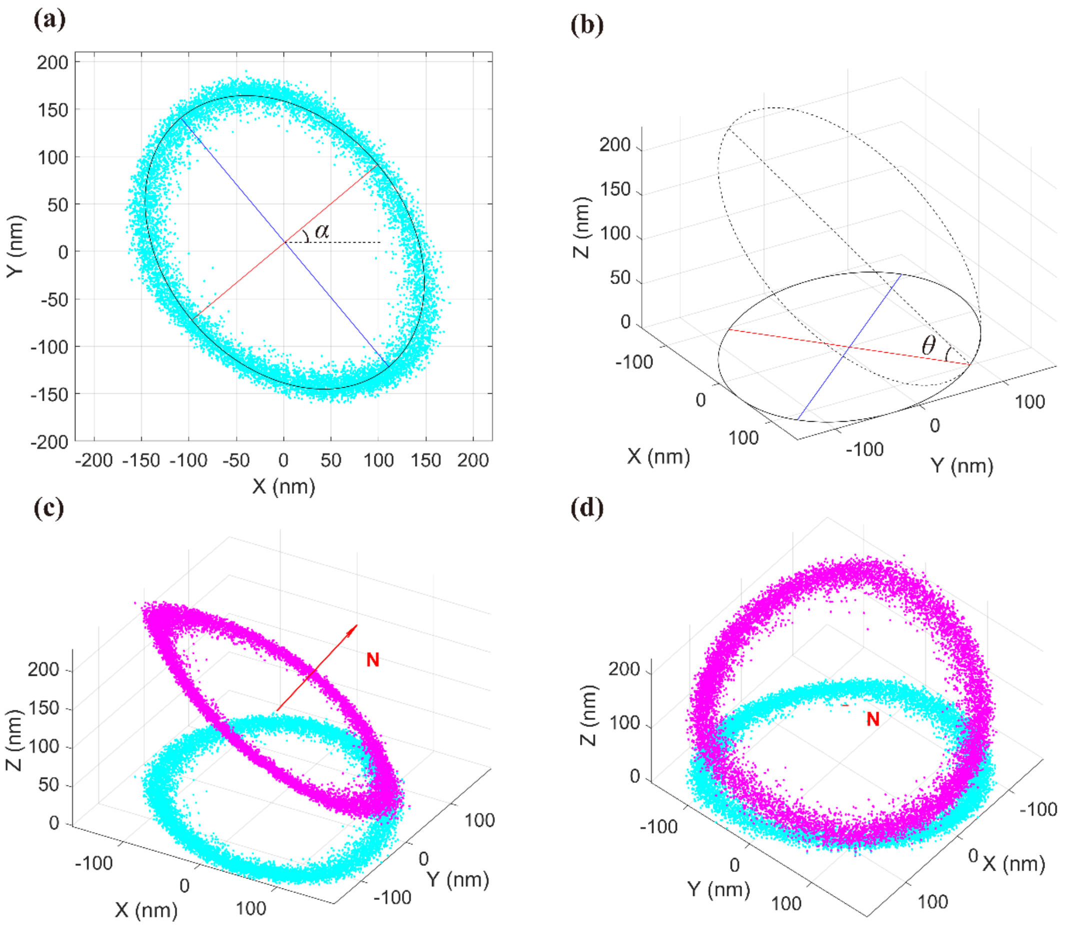


**Fig. S9.** Calculation of the normal vector of the bead rotation plane from an experimental trajectory. **(a)** Portion of the trajectory of a bead (the diameter of the bead is 1 μm) and its fitted ellipse (black line), as well as the major axis (blue line) and minor axis (red line) of the ellipse. The dashed black line is parallel to the X axis of the image coordinate system, and the angle between the minor axis and the X axis is $\alpha$. **(b)** A projection diagram, where a perfect circle (black dashed line) in 3d is projected onto the image focal plane as an ellipse (black solid line) and the cosine of inclination angle $\theta$ is the minor-to-major axis ratio. **(c)** The projected trajectory on the focal plane (cyan dots), the original trajectory in 3d on the rotation plane (magenta dots), and the normal vector of the rotation plane $\boldsymbol{N}$ (red arrow). **(d)** A view of (c) from the direction facing the normal vector $\boldsymbol{N}$.
